# hRBPome: a central repository of all known human RNA-binding proteins

**DOI:** 10.1101/269043

**Authors:** Pritha Ghosh, Pavalam Murugavel, Ramanathan Sowdhamini

## Abstract

**Background:** RNA-binding proteins (RBPs) have been extensively studied in humans over the past few years. Multiple reports have been made in the literature for genome-wide survey for this class of proteins in the human genome using different tools and techniques. Due to the inherent difference in the nature of the methods used in identifying human RBPs, as well as the diverse sources for data collection in each of these studies, there exists immense heterogeneity (including diversity in data formats) and poor intersection among the datasets available from the different studies.

**Description:** hRBPome is a comprehensive database of human RBPs known from six different studies. We have introduced considerable uniformity in the data, by mapping the various data formats reported by the different studies to gene names. This makes comparison across studies (datasets) easier than was possible before. We also provide confidence scores to each of these known RBPs, on the basis of their presence across studies.

**Conclusion:** This database presents a set of 837 high confidence RBP genes, identified in three or more resources, out of the six studies considered. Hence, it forms a “gold standard” for RBPs in the human genome. It also provides information for all the human RBPs from multiple resources, known to the best of our knowledge, on a common platform. The database can be accessed from the following URL: http://caps.ncbs.res.in/hrbpome

## Background

RNA-binding proteins (RBPs) are an important class of cellular proteins that are involved in various functions like stabilisation, protection, packaging, transport and mediating interactions with or act catalytically on RNA (cutting, unwinding, replicating, modifying, *etc*.) [1]. RBP-RNA associations are dynamic in nature and defines the rate of translation of mRNA cellular localisation, as well as the lifetime and processing of different kinds of RNA [2]. Our previous work has identified the complete RBP repertoire (or RBPome) in humans [3], but in the process of studying the same, we had realised the differences exist in the size of the human RBPome as reported by different groups [3–8]. To assemble all the known human RBPs, and their related information in one platform, we present hRBPome – a comprehensive database of human RBPs.

The six different studies (datasets) considered in hRBPome have been presented in **Table 1**. We have primarily focussed on highlighting the methods for identifying RBPs, adopted by each of these studies, as well as the start points for data collection (for example, human cell type for experimental studies or sequence database for computational studies), which lead to variation in the data formats reported. In this database, we have provided uniformity in the data, by mapping the various data formats reported by the different studies to gene names. This makes comparison across studies (datasets) easier than was possible before. The database can be accessed from the following URL: http://caps.ncbs.res.in/hrbpome

**Table 1:**
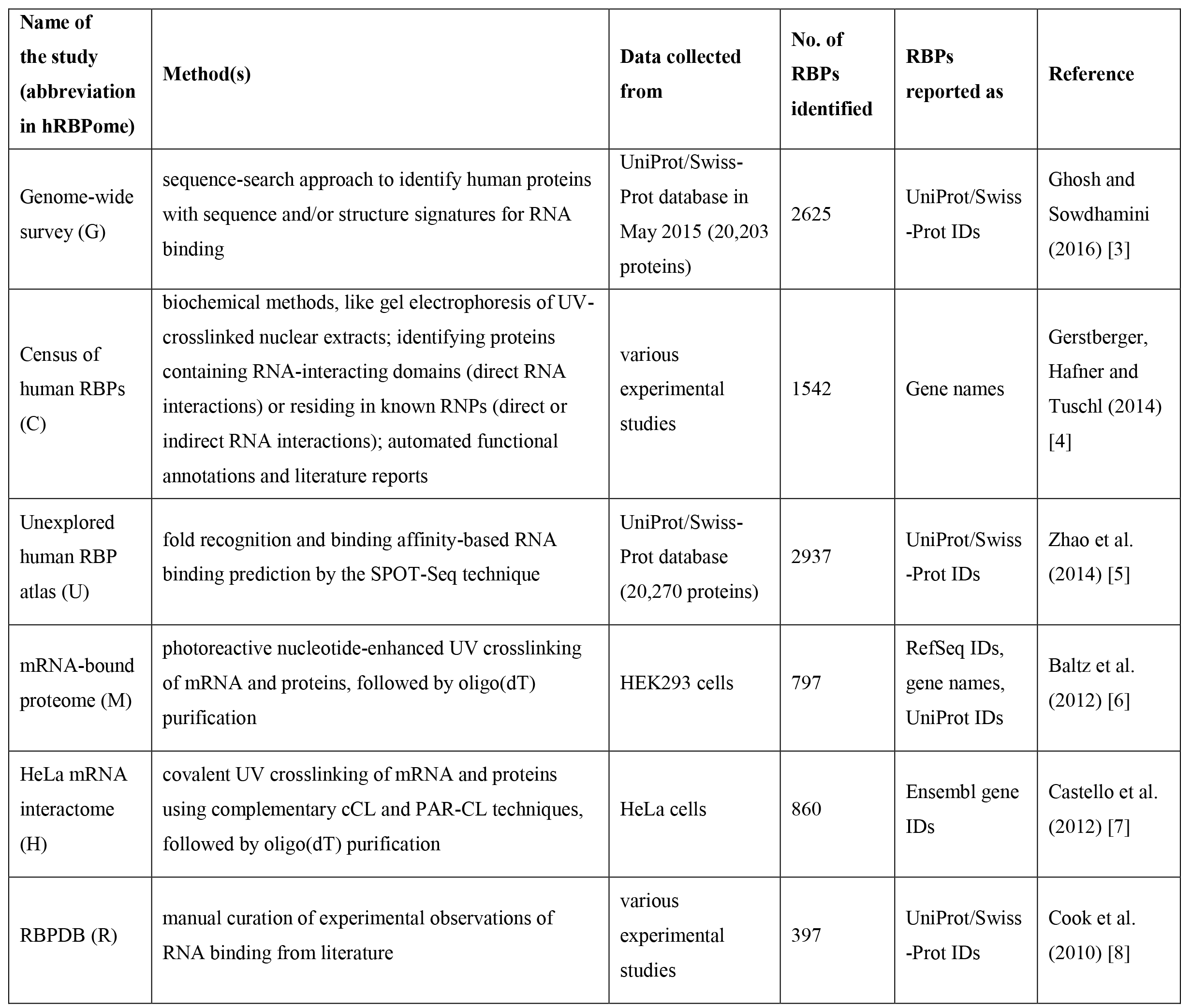
Details of the studies present in hRBPome. Various details of the studies (datasets) present in hRBPome have been tabulated here. We have primarily focussed on the methods adopted and the data formats in which the RBPs have been reported by each of these studies.

| Name of the study (abbreviation in hRBPome) | Method(s) | Data collected from | No. of RBPs identified | RBPs reported as | Reference |
| --- | --- | --- | --- | --- | --- |
| Genome-wide survey (G) | sequence-search approach to identify human proteins with sequence and/or structure signatures for RNA binding | UniProt/Swiss-Prot database in May 2015 (20,203 proteins) | 2625 | UniProt/Swiss-Prot IDs | Ghosh and Sowdhamini (2016) [3] |
| Census of human RBPs (C) | biochemical methods, like gel electrophoresis of UV-crosslinked nuclear extracts; identifying proteins containing RNA-interacting domains (direct RNA interactions) or residing in known RNPs (direct or indirect RNA interactions); automated functional annotations and literature reports | various experimental studies | 1542 | Gene names | Gerstberger, Hafner and Tuschl (2014) [4] |
| Unexplored human RBP atlas (U) | fold recognition and binding affinity-based RNA binding prediction by the SPOT-Seq technique | UniProt/Swiss-Prot database (20,270 proteins) | 2937 | UniProt/Swiss-Prot IDs | Zhao et al. (2014) [5] |
| mRNA-bound proteome (M) | photoreactive nucleotide-enhanced UV crosslinking of mRNA and proteins, followed by oligo(dT) purification | HEK293 cells | 797 | RefSeq IDs, gene names, UniProt IDs | Baltz et al. (2012) [6] |
| HeLa mRNA interactome (H) | covalent UV crosslinking of mRNA and proteins using complementary cCL and PAR-CL techniques, followed by oligo(dT) purification | HeLa cells | 860 | Ensembl gene IDs | Castello et al. (2012) [7] |
| RBPDB (R) | manual curation of experimental observations of RNA binding from literature | various experimental studies | 397 | UniProt/Swiss-Prot IDs | Cook et al. (2010) [8] |

The number of RBP genes that are common among the different studies (datasets) present in hRBPome have been represented as pairwise comparisons in **Figure 1**. There are approximately 47.5% common RBP genes among the M and the H datasets. The poor overlap among the other datasets have possibly arisen due the difference in the nature of identifying RBPs, as well as the use of diverse start points in each of these studies.

**Figure 1:**
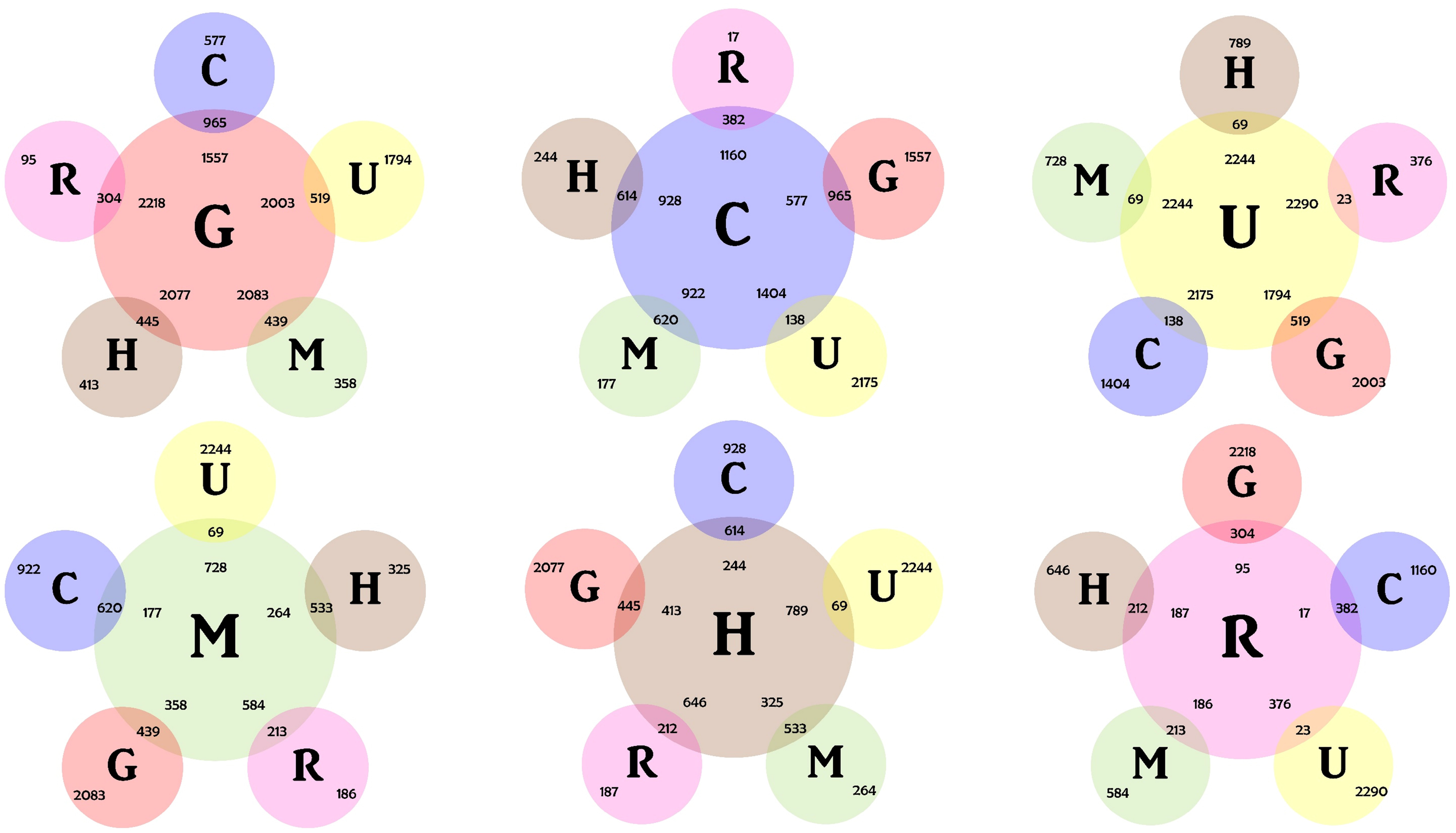
Comparative statistics of the different datasets present in hRBPome. The number of RBP genes that are common among the different studies (datasets) present in hRBPome have been represented as pairwise comparisons in this figure. In each panel, absolute numbers of common and unique RBP genes among two studies being compared, have been depicted. The abbreviations of the studies (datasets) are as follows: Genome-wide survey (G), Census of human RBPs (C), Unexplored human RBP atlas (U), mRNA-bound proteome (M), HeLa mRNA interactome (H) and RBPDB (R). There are approximately 47.5% common RBP genes among the M and the H datasets. The poor overlap among the other datasets have possibly arisen due the difference in the nature of identifying RBPs, as well as the use of diverse start points in each of these studies.

## Construction and content

The back-end data retrieval and manipulation logic of this database has been implemented using CGI-Perl and the database interface built on HTML5, CSS, JavaScript, Ajax and JQuery. The basic tables in the database have been organised as comma-separated text files, and converted to JSon format, for performance improvement through utilities.

The abbreviations of the studies (datasets) present in hRBPome are as follows: Genome-wide survey (G), Census of human RBPs (C), Unexplored human RBP atlas (U), mRNA-bound proteome (M), HeLa mRNA interactome (H) and RBPDB (R). The human RBPs reported by each of the six studies have been presented in this database, alongside a comparative account of all these studies. As seen from the penultimate column of Table 1, there are differences in the data formats (like, UniProt IDs, gene names etc.) in which the individual studies have reported human RBPs identified. To compare and contrast the RBPs reported by all the studies and also to provide a uniformity in the available data on human RBPs, ‘UniProt/Swiss-Prot IDs’ (from G, U and R) and ‘Ensembl gene IDs’ (from H) were mapped to ‘gene names’.

Further, the human RBP genes (*mapped* gene names from G, U, H and R, and the *reported* gene names from C and M) were compared across all the studies and this data is available in hRBPome for the users to browse. The genes that are present in three or more out of the six studies (G, U, H, R, C or M datasets), have been referred to as ‘high confidence RBP genes’ and have been assigned a confidence score of 3 or higher (highlighted in red) that equals the number of datasets in which that particular gene is present. The genes with a confidence score of less than 3 have been referred to as ‘low confidence RBP genes’. The users can browse through the entries present in a particular table and even the database as a whole, on the basis of keyword or ID searches (**Figure 2** and **Additional File 1**). hRBPome presents the human RBP gene names for the individual studies, cross-referenced to UniProt [9], RefSeq [10], Ensembl [11] and EMBL [12] databases. Further details about the protein can be obtained from the UniProt page, hyperlinked at the UniProt cross-reference.

Gene Ontology (GO) annotations (biological processes, molecular functions and cellular components) (**Additional File 2**) and disease associations of the RBPs (**Additional File 3**), if available, have also been provided in this database. The high confidence RBP genes have been highlighted in red font. The sequences of all the RBPs present in hRBPome have been made available for the users to download, in a study-wise manner. The sequences of all the high confidence RBPs are also available for download (**Additional File 4**). The database provides a link to RStrucFam, a webserver from our lab, which predicts RBPs and their cognate RNA partner(s) from mere sequence information [13].

All the data available to the users in the hRBPome have been organised in a tabular format. The records of these tables can be searched and/or sorted. The users also have a flexibility on the number of fields to be viewed at a time.

**Figure 2:**
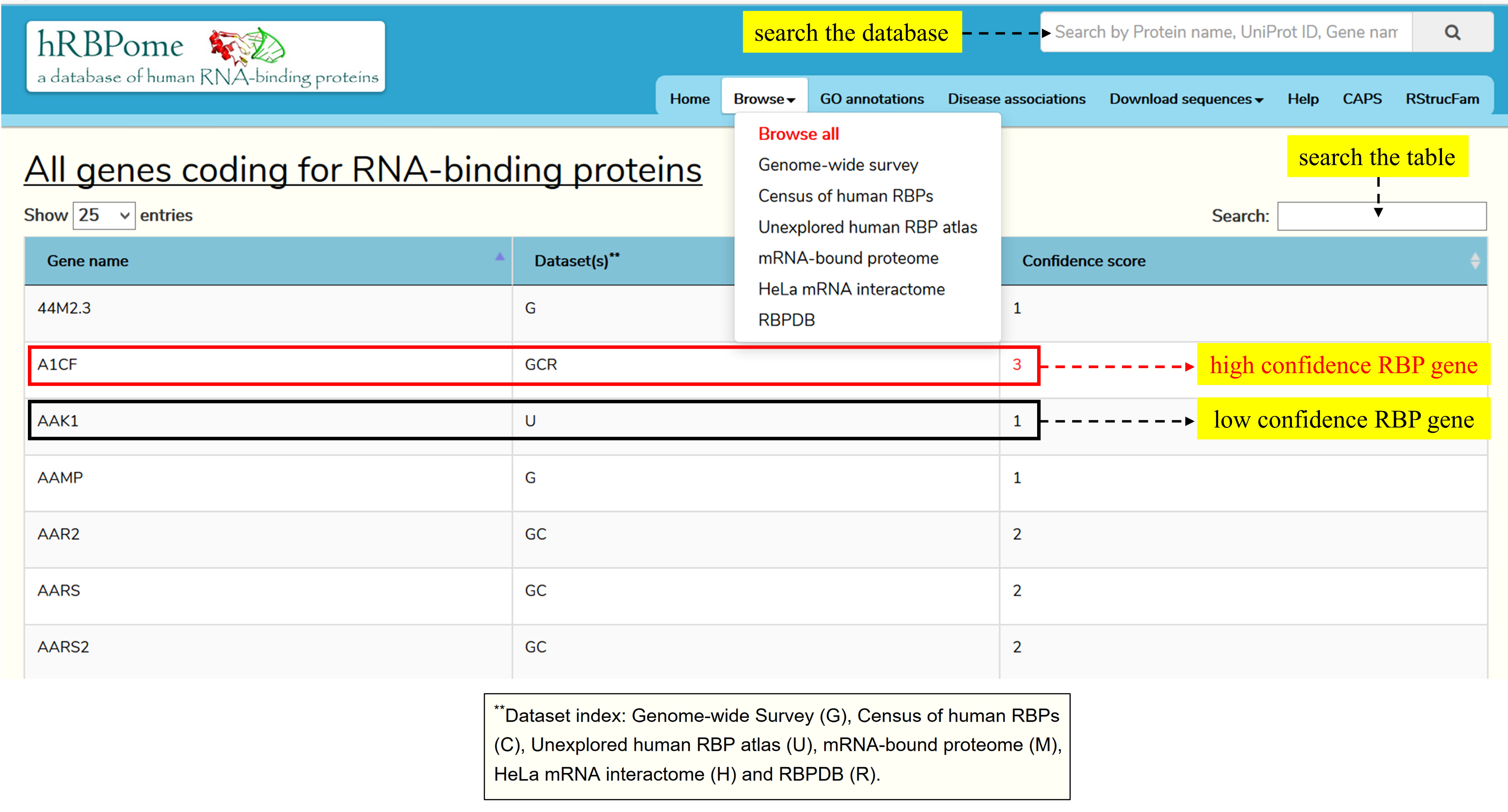
Browse through all the RBP genes present in hRBPome. The users can browse through all the genes coding for RBPs, which are present in hRBPome, by the virtue of being a part of one or more of the studies (datasets) considered in this database. The genes that are present in three or more out of the six studies (datasets), have been referred to as ‘high confidence RBP genes’ and have been assigned a confidence score of 3 or higher (highlighted in red) that equals the number of datasets in which that particular gene is present. The genes with a confidence score of less than 3 have been referred to as ‘low confidence RBP genes’. The users can browse through the entries present in a particular table and even the database as a whole, on the basis of keyword or ID searches.

## Utility and discussion

The hRBPome database described here, to the best of our knowledge, is the first of its kind attempt, to bring all known human RBPs in one platform. The data presented in this database also resolves differences arising out of the nature of tools and techniques used or that of the data presented, in the various existing studies for the identification of human RBPs. A comparison of human RBPs from multiple studies, hRBPome also annotates ~800 high confidence RBP genes. The entries present in this database are cross-referenced to various other protein sequence databases.

In 2014, Gerstberger and co-workers had proposed a set of 554 mRNA-binding protein (mRBP) genes as the “gold standard” [4]. Our current study provides a new gold standard of 837 high-confidence RBP genes, and includes not only genes encoding mRBPs, but also those encoding proteins binding to all other known RNA kinds. 499 RBP genes are common among the old and the new gold standards, whereas 55 and 338 genes are unique to the old and new gold standards, respectively (**Additional File 5**). In short, this database is a central repository for all human RBPs known till date and information related to these proteins.

### Conclusions

In the past few years, multiple research groups have sought to understand the complete RBP repertoire of humans. Due to the employment of a variety of different tools, these studies have presented non-overlapping datasets of human RBPs. There has been little realisation that differences could exist due to the choice of search techniques and thresholds placed. Furthermore, some techniques could also identify “indirect interactions” i.e. proteins that are part of assemblies and not directly interacting with RNA. Likewise, few techniques may identify DNA-binding proteins or proteins that can bind to both DNA- and RNA.

Though it is important to identify those proteins that are reported by multiple such studies, it is also interesting to note the union of these sets. The hRBPome database presents a comparative account of all known human RBPs, which provides objectivity to the meta integration, thereby data become easily available to users than ever before. Subsequent updates of the database will be targeted towards integration of more features of RBPs than currently available.

### List of abbreviations

C: Census of human RBPs
cCL: Conventional UV crosslinking
G: Genome-wide survey
GO: Gene Ontology
H: HeLa mRNA interactome
M: mRNA-bound proteome
mRBP: mRNA-binding protein
PAR-CL: Photoactivatable-ribonucleoside-enhanced crosslinking
R: RBPDB
RBP: RNA-binding protein
RNP: Ribonucleoprotein
U: Unexplored human RBP atlas
UV: Ultraviolet

## Declarations

All authors have gone through the manuscript and contents of this article have not been published elsewhere.

### Ethics approval and consent to participate

Not applicable

### Consent for publication

Not applicable

### Availability of data and material

The database can be accessed at http://caps.ncbs.res.in/hrbpome/

### Authors’ contributions

RS conceptualised the project. Curation of the database was performed by PG and PM. PG wrote the first draft of the manuscript and RS improved upon it. All the authors read and approved the final version of the manuscript.

## Acknowledgments

The authors acknowledge NCBS (TIFR) for infrastructural facilities.

## Additional Files

### Additional File 1: RBP genes present in individual studies

The users can browse through each of the studies (datasets) present in hRBPome. This has been exemplified here with our work on the genome-wide survey of human RBPs. The genes encoding for RBPs, as well as their cross-references from different databases like UniProt, RefSeq, Ensembl and EMBL have been provided. The high confidence RBP genes have been highlighted in red font.

### Additional File 2: GO annotations

GO annotations (biological processes, molecular functions and cellular components) of the RBPs, if available, have been provided in hRBPome. The high confidence RBP genes have been highlighted in red font.

### Additional File 3: Disease associations

Disease associations of the RBPs, if known, have been provided in hRBPome. The high confidence RBP genes have been highlighted in red font.

### Additional File 4: User downloads

The sequences of all the RBPs present in hRBPome have been made available for the users to download, in a study-wise manner. The sequences of all the high confidence RBPs are also available for download. The database provides a link to RStrucFam, a webserver from our lab, which predicts RBPs and their cognate RNA partner(s) from mere sequence information.

